# Nutrient deposition enhances post-fire survival in non-N-fixing savanna tree seedlings

**DOI:** 10.1101/444372

**Authors:** Varun Varma, Mahesh Sankaran

## Abstract

Nutrient deposition can modify plant growth rates and potentially alter the susceptibility of plants to disturbance events, while also influencing properties of disturbance regimes. In mixed tree-grass ecosystems, such as savannas and tropical dry forests, tree seedling growth rates strongly influence the ability of seedlings to survive fire (i.e. post-fire seedling survival), and hence, vegetation structure and tree community composition. However the effects of nutrient deposition on the susceptibility of recruiting trees to fire are poorly quantified. In a field experiment, seedlings of multiple N-fixing and non-N-fixing tropical dry forest tree species were exposed to nitrogen (N) and phosphorus (P) fertilisation, and fire. We quantified nutrient-mediated changes in a) mean seedling growth rates; b) growth rates of the fastest growing individuals and c) post-fire seedling survival. N-fixers had substantially higher baseline post-fire seedling survival, that was unaffected by nutrient addition. Fertilisation, especially with N, increased post-fire survival probabilities in non-N-fixers by increasing the growth rates of the fastest growing individuals. These results suggest that fertilisation can lead to an increase in the relative abundance of non-N-fixers in the resprout community, and thereby, alter the community composition of tropical savanna and dry forest tree communities in the long-term.

## Introduction

Increased atmospheric deposition of nitrogen (N) and phosphorus (P) as a consequence of human activities is an important global change driver (Vitousek 1994; Vitousek et al. 1997; Sala et al. 2000; Falkowski et al. 2000; Galloway et al. 2004, 2008; Phoenix et al. 2006, 2012; Dentener et al. 2006; Mahowald et al. 2008; Du and Liu 2014; Vet et al. 2014) that has been shown to modify vegetation communities in a number of ecosystems (Elser et al. 2007; Xia and Wan 2008; Bobbink et al. 2010). Numerous studies have demonstrated that increased availability of these essential plant nutrients can differentially affect plant growth amongst members of a community, bringing about compositional changes in vegetation assemblages (Huante et al. 1995a, b; Zavaleta et al. 2003; Khurana and Singh 2004; Stevens et al. 2004, 2006, 2010; Wassen et al. 2005; Elser et al. 2007; Clark and Tilman 2008; Xia and Wan 2008; Tripathi and Raghubanshi 2014; Verma et al. 2014; Powers et al. 2015). In addition, increased nutrient availability can also indirectly modify plant communities by affecting properties of local disturbance regimes – which play a ubiquitous role in shaping ecosystems across the globe – and by modifying the susceptibility of plant species to disturbance events (Heil and Diemont 1983; Bobbink and Lamers 2002; Strengbom et al. 2002; Bobbink et al. 2010; Barbosa et al. 2014b; Varma et al. 2018). To assess the net effect of nutrient deposition on vegetation communities, we need to therefore consider both direct nutrient-mediated effects on plant growth, as well as indirect effects that arise through alterations in disturbance characteristics and the differential susceptibility of plant species to these altered disturbance regimes.

Mixed tree-grass ecosystems, such as savannas and seasonally dry tropical forests with grassy understories, are globally important biomes (Murphy and Lugo 1986; Field et al. 1998; Powers and Tiffin 2010; McShea et al. 2011), covering approximately 20% of the Earth’s terrestrial surface (Scholes and Archer 1997). However, as with tropical ecosystems in general (Matson et al. 1999; Siddique et al. 2010; Bobbink et al. 2010; Lu et al. 2010), the effects of atmospheric nutrient deposition on the dynamics of mixed tree-grass ecosystems are relatively poorly investigated (Powers et al. 2015). Vegetation in these systems is characterised by a spatially discontinuous tree canopy, interspersed within a continuous layer of C 4 grasses (Scholes and Archer 1997; Bond et al. 2003a; House et al. 2003; Bond 2008; Lehmann et al. 2009b; Edwards et al. 2010; Bond and Parr 2010; Ratnam et al. 2011). The availability of resources such as water and nutrients, as well as disturbances, such as fire and herbivory, strongly influence the extent of woody cover and tree species diversity in these communities (Sankaran et al. 2004, 2005, 2008; Lehmann et al. 2009a, 2014; Staver et al. 2009; Dantas et al. 2015). They are also amongst the most frequently burned biomes in the world (Dwyer et al. 1998, 2000; Barbosa et al. 1999; Tansey et al. 2004; Bond et al. 2005), and in the wetter regions of these biomes (e.g. in Africa, where annual precipitation > 650 mm), grass-fuelled fires are the single most important constraint to the recruitment of trees into the adult canopy layer (Higgins et al. 2000; Sankaran et al. 2005; Bond 2008).

In the long-term, nutrient deposition has the potential to alter tree canopy cover in mixed tree-grass ecosystems, both directly, through their effects on tree seedling growth, as well as indirectly via effects on grass growth, which in turn can modify the strength of tree-grass competition and the nature of fire regimes. Tree seedlings in general demonstrate enhanced growth with nutrient availability (Huante et al. 1995a; Khurana and Singh 2004; Vadigi and Ward 2013; Varma et al. 2018) and may therefore be expected to show increased post-fire survival because larger seedlings survive fires better (Trollope 1984; Gignoux et al. 1997; Hoffmann and Solbrig 2003; Bond 2008; Lawes et al. 2011; Wakeling et al. 2011). However, nutrient-mediated increases in seedling growth are also typically associated with reduced relative investment in root biomass, as well as root carbohydrates (Tilman 1988; Khurana and Singh 2004; Knox and Clarke 2005; Hermans et al. 2006; Wang et al. 2015; Varma et al. 2018), which in turn could negatively influence post-fire seedling survival and resprouting ability (Hoffmann et al. 2000; Bell 2001; Bond et al. 2003b; Lamont and Wiens 2003; Vesk and Westoby 2004; Clarke and Knox 2009; Wigley et al. 2009; Clarke et al. 2013). In addition, increases in grass productivity with fertilisation (Fynn and O’Connor 2005; Lee et al. 2010) has the potential to negatively affect tree seedling growth and survival, both as a result of increased grass competition (Higgins et al. 2000; Kraaij and Ward 2006; Cramer et al. 2010; Kambatuku et al. 2012; Cramer and Bond 2013; Vadigi and Ward 2013, 2014), as well as more intense and frequent grass-fuelled fires (Williams et al. 1999; Higgins et al. 2000; Ryan and Williams 2011; Hoffmann et al. 2012).

In addition to influencing structure, N and P deposition can also potentially lead to changes in the functional composition of tree communities in mixed tree-grass ecosystems if they have differential effects on the growth and survival of dominant tree functional types in the system, i.e. N-fixers and non-N-fixers. N-fixers and non-N-fixers have contrasting nutrient demands and acquisition strategies (Vitousek et al. 2002, 2013; Pearson and Vitousek 2002) which can result in differing growth responses of these functional groups to nutrient enrichment (Khurana and Singh 2004; Cramer et al. 2007; Barbosa et al. 2014a; Tripathi and Raghubanshi 2014; Varma et al. 2018). Furthermore, differences between functional groups in their ability to withstand grass competition (Cramer et al. 2007, 2010), and in biomass allocation patterns to above-versus below-ground organs and root carbohydrate reserves (Khurana and Singh 2004; Tripathi and Raghubanshi 2014; Varma et al. 2018) can result in differing post-fire resprouting and survival with N and P fertilisation.

Ultimately, the long-term effects of nutrient deposition on the structure and composition of mixed tree-grass systems will depend on the relative strengths of these direct and indirect nutrient-mediated effects. Although a few previous studies have investigated how nutrient availability and fire influence the growth and survival of N-fixing and non-N-fixing species (Kraaij and Ward 2006; Higgins et al. 2007; Cramer et al. 2007, 2010, 2012; Kambatuku et al. 2012; Vadigi and Ward 2012, 2013, 2014; Cramer and Bond 2013; Barbosa et al. 2014a, b), particularly in African savannas, our understanding of the individual and interactive effects of nutrients and fire on tree demography in these systems remains incomplete. Few studies have looked at the effects of N and P separately, and fewer still, in the context of Asian savannas.

Here, we report results from a field experiment that investigated the effects of N and P fertilisation and fire on seedling growth and survival of multiple N-fixing and non-N-fixing tropical savanna and dry forest tree species in India. The main aim of this experiment was to determine the extent to which projected increases in anthropogenic N and P deposition are likely to affect tree seedling establishment in tropical savannas and dry forests of the Indian subcontinent, and thereby, potentially alter tree demography and functional composition of the region. More specifically, our objectives were to quantify the effects of N and P fertilisation on 1) seedling growth rates, including both mean growth rates as well as growth rates of the fastest growing individuals (Wakeling et al. 2011) of N-fixing and non-N-fixing species, in the presence of grass competition, 2) grass biomass production, and thus, fuel load accumulation, and 3) post-fire seedling survival of N-fixing and non-N-fixing species.

## Methods

The experiment was conducted on 0.5 ha of abandoned farm land in the village of Hosur, located in Mysore district of the southern Indian state of Karnataka. The region receives approximately 800 mm of rainfall annually, primarily from the south-west monsoon between June and October (Indian Meteorological Department – http://www.imd.gov.in). This is followed by a dry season from December to April. Fires are typically observed between February and April when average day-time temperatures range from 31°C to 35°C.

Between August and October, 2012, prior to the start of the experiment, the top 30 cm of soil was homogenised with two rounds of ploughing, and the experimental area was fenced to exclude livestock and human disturbance. Forty eight experimental plots were subsequently established (8m X 8m) by placing non-leaching plastic lining up to a depth of 0.5 m around them to prevent the movement of added nutrients between plots. Grasses were allowed to naturally regenerate in the experimental plots.

Eight commonly occurring tropical dry forest tree species (Puyravaud et al. 1994; Sagar and Singh 2004; Kumar and Shahabuddin 2005; Kodandapani et al. 2008) were selected for the experiment which included four N-fixers (*Acacia ferruginea* DC., *Albizia amara* (Roxb.) B.Biovin., *Albizia lebbeck* (L.) Benth. and *Dalbergia latifolia* (Roxb.)) and four non-N-fixers (*Lagerstroemia speciosa* (L.), *Sapindus emarginatus* Vahl., *Terminalia arjuna* (Roxb. ex DC.) and *Ziziphus jujuba* Mill.). All N-fixers used in this experiment have previously been shown to nodulate (Germplasm Resource Information Network – http://www.ars-grin.gov/; also see Varma et al. 2018). Seedlings of the eight species were obtained from the Foundation for Revitalisation of Local Health Traditions (FRLHT), Bangalore and transported to the field site six weeks after germination. In November 2012, after a two-week acclimation period, 68 seedlings across all eight species were planted into each treatment plot. Plants were placed at random within a grid with a minimum spacing of 80 cm between adjacent plants. The number of individuals per species varied within plots (*A. ferruginea* = 10, *A. amara* = 6, *A. lebbeck* = 5, *D. latifolia* = 10, *L. speciosa* =10, *S. emarginatus* = 8, *T. arjuna* = 9, *Z. jujuba* = 10). However, each plot contained identical numbers of individuals of each species at the start of the experiment. To aid establishment and reduce seedling mortality, additional water was supplied to seedlings manually and using a sprinkler system from November 2012 to June 2013.

Twelve treatment plots were randomly assigned to one of four nutrient treatments – control (no added nutrients), N+ (2 g N.m^−2^.y^−1^), P+ (0.2 g P.m^−2^.y^−1^) and NP+ (2 g N.m^−2^.y^−1^ and 0.2 g P.m^−2^.y^−1^). The N and P addition levels used represent conservative estimates of atmospheric N (Dentener et al. 2006) and P (Mahowald et al. 2008) deposition expected by 2030 over southern India. Nutrients were evenly applied to each treatment plot as solutions of urea and single superphosphate for N and P, respectively. Nutrient additions were carried out in two applications in November-December, 2012, three applications in July-August, 2013 and three applications in July-August, 2014. Seedling height (using a tape measure) and basal stem diameter (measured 2 cm above the ground using a vernier calliper) were measured for each individual one-week after transplant in November 2012 (H_1_ and D_1_ hereafter for height and stem diameter, respectively) and again 15 months later in February 2014 (H_2_ and D_2_ hereafter for height and stem diameter, respectively). Seedling relative growth rates for stem height and diameter were calculated as (H_2_ – H_1_)/H_1_ and (D_2_ – D_1_)/D_1_, respectively. Additionally, in February 2014, herbaceous biomass from three 10 cm x 20 cm quadrats (separated into grass and forb biomass) was harvested from each of the 48 experimental plots to quantify fuel load.

Coinciding with the peak of the dry season in February-March 2014, six plots from each nutrient treatment level (total of 24 plots) were selected at random for the fire treatment. Prior to burning, all above-ground biomass in plots (excluding the transplanted seedlings) was clipped and placed back within the plots to facilitate even drying of the fuel layer and prevent fires from spreading to adjacent plots. Flame heights during burning were observed to be well in excess of the heights of most seedlings, suggesting that pre-burn clipping of the grass layer was unlikely to have significantly modified the effect of fire on seedlings. Seedling survival was quantified at the end of the wet season following burning (September 2014). Seedlings that showed no live above-ground biomass, i.e. had not resprouted, were scored as dead.

### Data analysis

The effects of N and P addition on pre-burn relative growth rates of tree seedlings were evaluated using linear mixed effects models, with nutrient treatment and functional group included as fixed effects. Species identity was used a random factor within these models, as the focus of the analysis was to detect functional group level generalisations, while accounting for intrinsic species-specific differences in relative growth rates. In addition, data from plants in the control treatment were used to quantify baseline differences in the growth rates of N-fixers and non-N-fixers. Growth rates for seedlings in the control treatment were modelled as a function of initial size and plant functional group, with species identity included as a random factor.

A bootstrapping method was used to quantify the effect of nutrients on growth rates of the fastest growing seedlings. Growth rates of all individuals were resampled with replacement (100 iterations) for each species-treatment combination separately. At each iteration, the log response ratio (LRR) between the 75^th^ percentile of growth rates in the nutrient treatment and that of the control treatment were calculated. Changes in the growth rates of the fastest growing individuals with fertilisation were represented by the mean LRR and associated 95% confidence intervals (CIs), i.e. the LRR represents the change in 75^th^ percentile growth rate of a species with fertilisation and the 95% CIs around the LRR captures the variation around the response. In addition, mean 75^th^ percentile growth rates and 95% CIs in the control treatment for each species were also reported. The effects of nutrient treatments on grass, forb and total herbaceous biomass were analysed using separate linear mixed effects models, where plot identity was used as a random factor.

The effect of N and P fertilisation on seedling mortality following fire was analysed using a generalised linear mixed effects model with a binomial error distribution. The analysis allowed for a three-way interaction between nutrient treatment, fire treatment (unburned and burned) and plant functional group. Species identity was included as a random factor. Additionally, we assessed the effect of plant size (H 2 and D2 – measured before the application of fire) and nutrient treatment on the probability of seedling survival. To avoid an excessively complicated analysis framework and to facilitate a clearer presentation of results, this analysis was carried out for each species separately using generalised linear models with binomial error distributions. This procedure involved the construction of two models for each species. The first model, which assessed the overall effect of seedling size on post-fire survival, pooled data across nutrient treatments for a given species and included seedling survival as a binary response variable, and seedling size as a predictor. The second model assessed the effects of nutrient addition on seedling survival and included nutrient treatments and plant size as predictors. This second model was subjected to a backward stepwise elimination of terms to arrive at a minimum acceptable model.

All statistical analyses were carried out in R (R Core Team 2014). We used the *lme4* package (Bates et al. 2014) for analyses that involved linear and generalised linear mixed effects models. The statistical significance of parameter estimates of the mixed models were computed using Satterthwaite’s approximation of degrees of freedom implemented within the *lmerTest* package (Kuznetova et al. 2014). However, we place equal emphasis on the magnitude of effects and do not rely solely on computed *P* values to interpret results.

## Results

N-fixers had marginally higher growth rates compared to non-N-fixers in the control treatment (Fig 1; *P* = 0.054). Growth rates decreased as a function of initial plant size, but patterns did not differ between functional groups (initial plant size x functional group: NS). Averaging across all nutrient treatments, the rate of stem height increments in N-fixers was 3.3 times that observed in non-N-fixers in the 15 months between measurements, but growth rates in both functional groups were very variable. Nutrient addition did not affect mean growth rates in either functional group (i.e. nutrient treatment x functional group: NS, nutrient treatment: NS).

**Fig 1.**
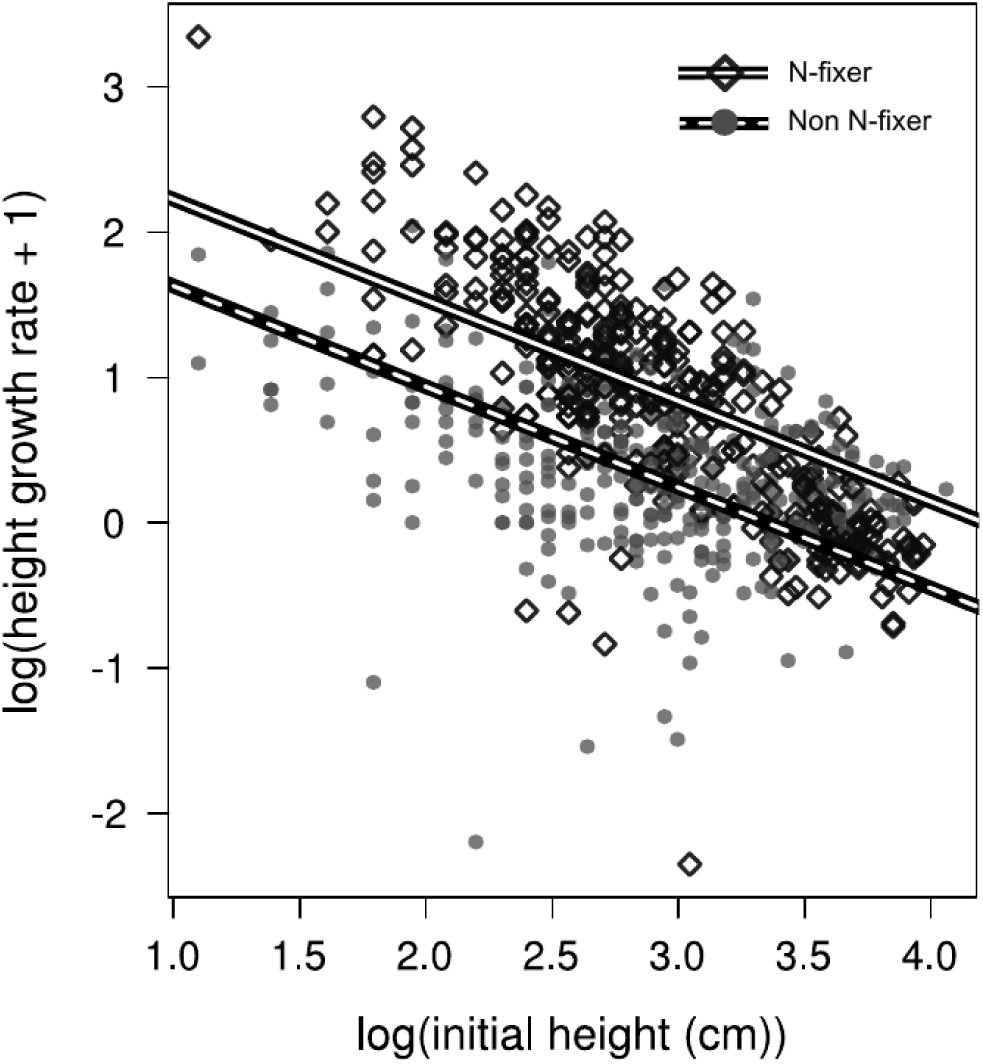
Relationship between relative height growth rate and initial seedling height in the control treatment (no nutrient addition). N fixers show marginally faster mean growth rates compared to non-N-fixers (*P* = 0.054)

N-fixers in general had greater growth rates amongst the fastest growing individuals in terms of both stem height (Fig 2a) and diameter growth (Fig 2b). The only exception to this was *A. ferruginea* which had very slow height growth rates (Fig 2a). Fertilisation resulted in large changes in the growth rates of the fastest growing individuals. Amongst N-fixers, nutrient addition largely increased stem height growth rates (positive LRRs where CIs did not overlap with zero), except in *D. latifolia* which showed a small decline (Fig 2a). We also note that the large increases observed for *A. ferruginea* could be an artefact of slow growth rates for this species in general (Fig 2a). With respect to stem diameter growth rates, N-fixers showed a more mixed response to fertilisation across species. However, non-N-fixers showed consistent increases in stem diameter growth rates (Fig 2b). As a consequence, in all four non-N-fixing species, greater stem diameters were observed in nutrient addition treatments, especially with N addition, compared to the maximum stem diameter achieved in the control treatment for each species (Fig 3).

**Fig 2.**
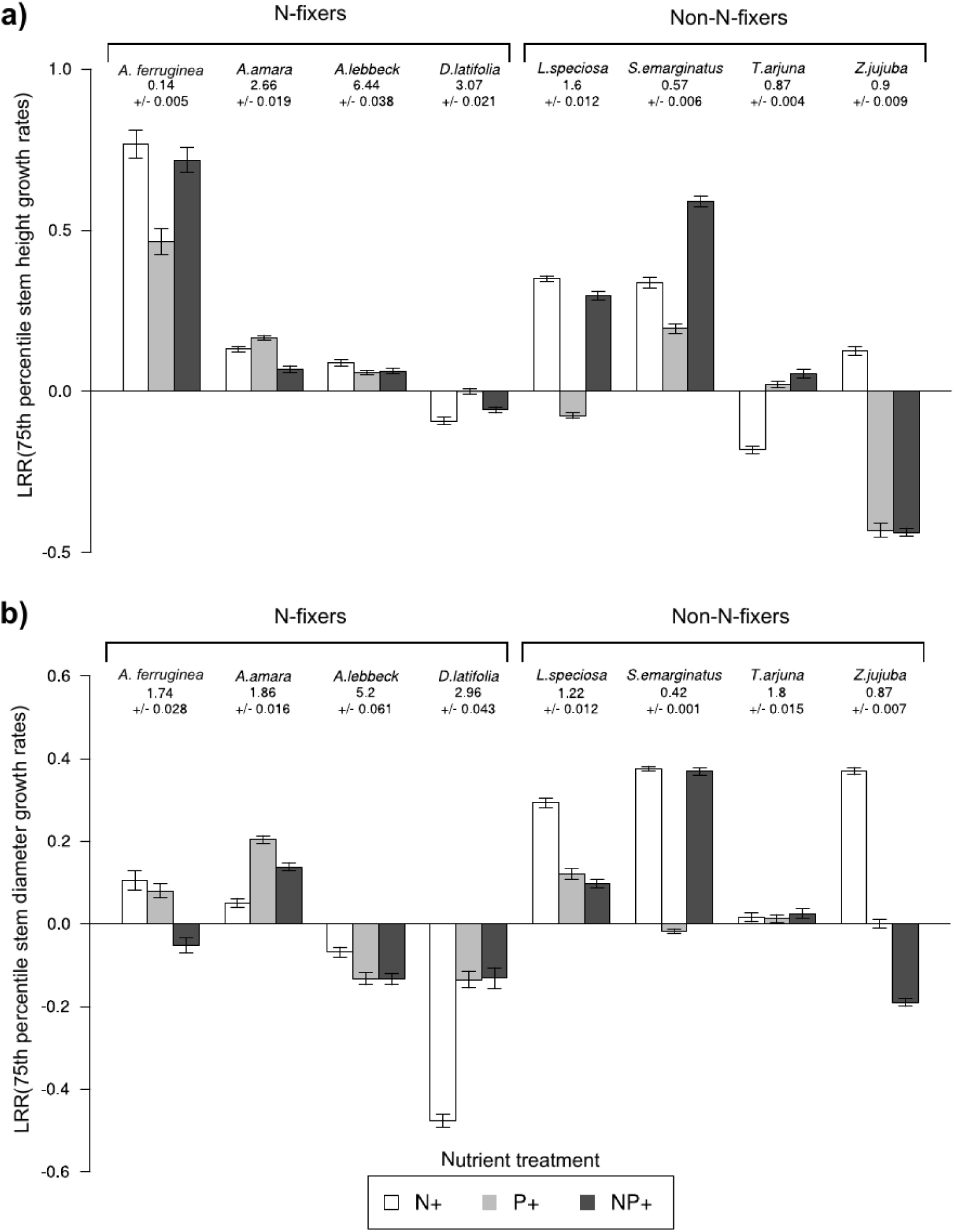
Bootstrapped log response ratios (LRR) of the 75^th^ percentile growth rates for stem height (a) and stem diameter (b). Error bars indicate 95% confidence intervals. Values quotes below species names are bootstrapped mean 75^th^ percentile growth rates observed in the control treatment (no nutrient addition) for the respective species and associated 95% confidence intervals.

**Fig 3.**
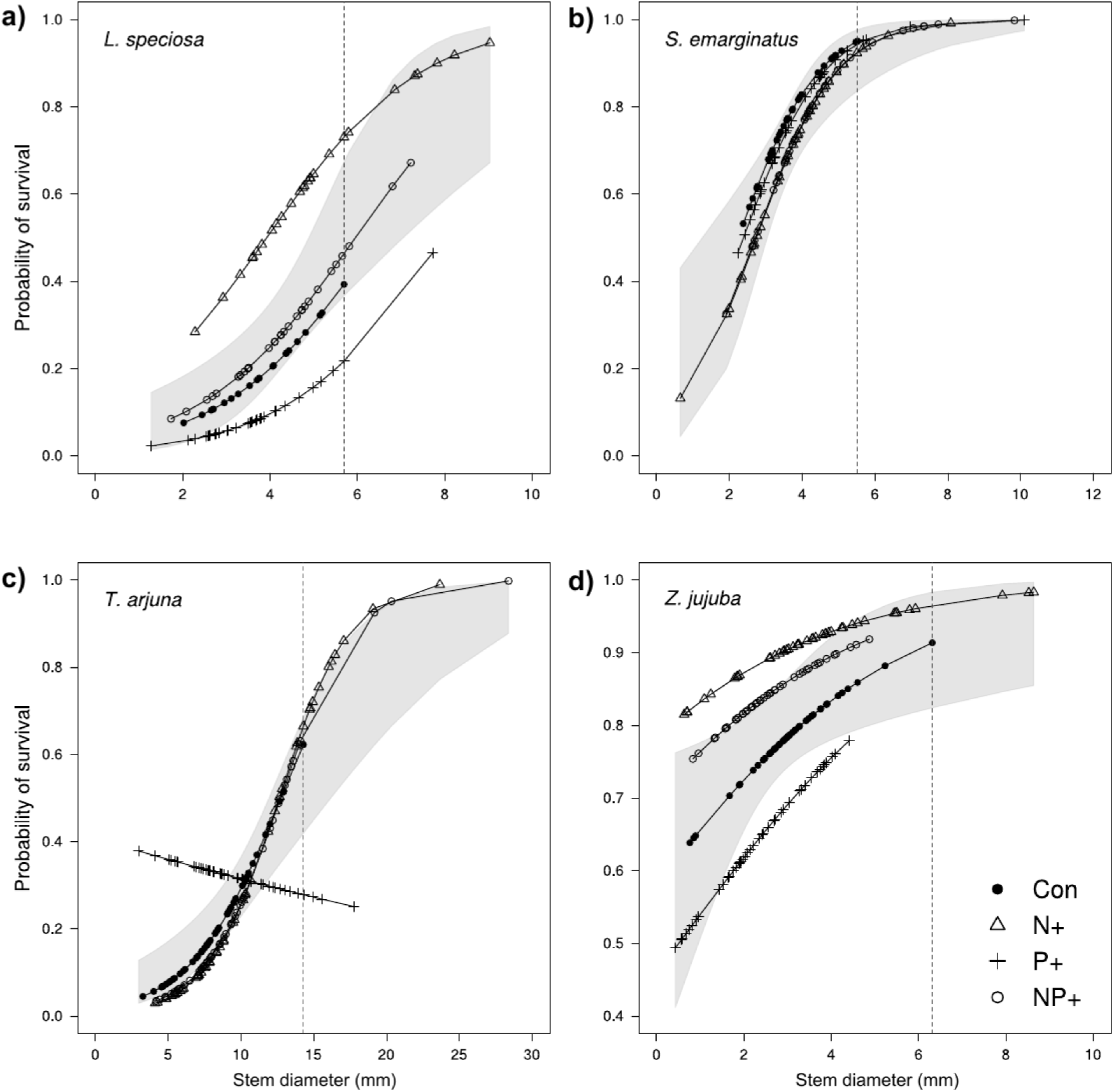
Relationship between seedling pre-burn stem diameter and fitted probability of survival when burned (generalised linear model with binomial error distributions) for each non-N-fixing species – *L. speciosa* (a), *S. emarginatus* (b), *T. arjuna* (c) and *Z. jujuba* (d). Shaded regions represent the 95% confidence bounds of the relationship between stem diameter and survival probabilities when all nutrient treatments are pooled. The vertical dotted lines indicate the maximum stem diameter observed in the control treatment (no nutrient addition) for each species.

Seedling survival was in general very high in N-fixers. In the unburned treatment, 99% of N-fixers survived compared to 87% in the case of non-N-fixers (Fig 4a-b; *P* < 0.001). Amongst interactions included in the model, the three-way interaction between functional group, nutrient treatment and fire treatment was not significant, and only the two-way interaction between functional group and fire treatment was a significant predictor of seedling survival (*P* < 0.001; *df*_1, 384_). Burning had no effect on N-fixer survival which remained high (99% resprouted following burning; Fig 4a), but significantly reduced survival in non-N-fixers with only 51% resprouting following burning (Fig 4b; *P* < 0.001). Of the 1364 seedlings exposed to fire, only four individuals, all of which were N-fixers, reached sufficient height (> 140 cm) to escape top-kill. Thus, fire-mediated mortality was almost exclusively restricted to non-N-fixers. Nutrient treatments did not affect seedling survival in either functional group (Fig 4a, b), though high variation in survival was observed in non-N-fixers exposed to N addition (Fig 4b). Grass and forb biomass production was not affected by fertilisation. Therefore, nutrient deposition at application rates used in this experiment did not influence fuel loads, and hence, fire intensity.

**Fig 4.**
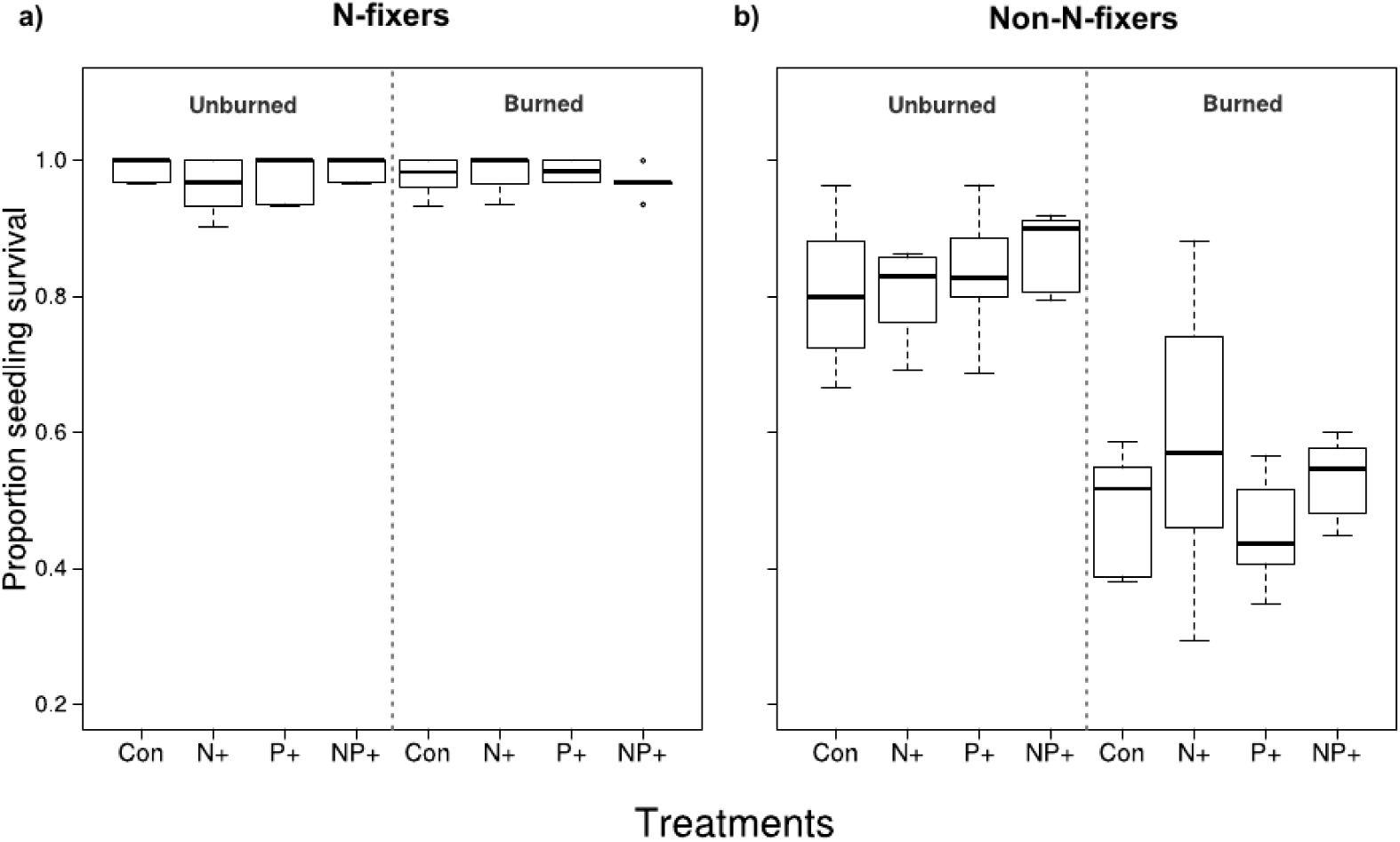
Observed seedling survival in unburned and burned treatments for N-fixers (a) and non-N-fixers (b).

In non-N-fixing species, increasing stem diameter and height (data not shown) significantly increased probability of post-fire seedling survival (*P* < 0.001 for all species; Fig 3a-d). The only exception to this relationship was observed in *T. arjuna* seedlings exposed to P addition, where survival declined marginally with increasing stem diameter (Fig 3c). In non-N-fixers, nutrient addition, especially N, increased stem diameter growth rates of the fastest growing individuals (Fig 2b) and resulted in seedlings reaching larger stem diameters in fertilised plots compared to their respective controls (Fig 3a-d). In turn, these larger seedlings had higher survival probabilities. In contrast, in the case of N-fixers, there was no relationship evident between post-fire survival and plant size (i.e. stem height and diameter) given that virtually all individuals survived fire.

## Discussion

N-fixing tropical dry forest seedlings showed faster growth rates compared to non-N-fixers. N-fixers also demonstrated a substantially greater ability to survive fires. Nutrient addition did not affect mean seedling growth rates in either functional group in this study. However, fertilisation had a strong influence on the growth rates of the fastest growing seedlings in both functional groups. Consistent increases in stem diameter growth rates of the fastest growing individuals of non-N-fixing species allowed these seedlings to attain larger stem diameters, which in turn, increased their survival probabilities following fires. Although N-fixing species were largely unaffected by either fire or nutrient addition, with virtually all individuals surviving, the increased growth and survival probabilities of the fastest growing non-N-fixers with nutrient addition suggests that non-N-fixers may benefit from increased nutrient deposition in the future.

In the control treatment, i.e. with no added nutrients, N-fixers showed marginally higher relative growth rates compared to non-N-fixers. Faster growth rates in N-fixers could be attributed to greater foliar N content (Khurana and Singh 2004; Wright et al. 2004; Powers and Tiffin 2010; Orwin et al. 2010; Reich 2014) afforded by their association with N-fixing bacteria in root nodules. As a consequence, N-fixers were taller at the time of burning and showed higher post-fire survival compared to non-N-fixers. N and P addition in general did not affect mean seedling growth rates in both functional groups. However, fertilisation did result in changes in growth rates of the fastest growing individuals of a species, especially for non-N-fixers with N addition. Wakeling et al. (2011) argued that growth rates of the fastest growing individuals (the fastest 5%), as opposed to mean growth rates, were better predictors of the number of individuals escaping fires, and consequently, of observed tree densities in an African savanna (Wakeling et al. 2011). Hence, non-N-fixers are likely to benefit from enhanced in N deposition through consistent increases in the growth rates of the fastest growing individuals and resulting increases in post-fire survival.

In contrast to many previous studies (Fynn and O’Connor 2005; Craine et al. 2008; Kambatuku et al. 2012; Cramer et al. 2012; Vadigi and Ward 2012; Verma et al. 2014), fertilisation did not influence mean seedling growth rates and herbaceous biomass production in this experiment. Studies which have reported increases in herbaceous production and tree seedling growth rates previously, have used N application rates that were between three and six times higher than that used here (Kambatuku et al. 2012; Cramer et al. 2012; Verma et al. 2014). Our results are, however, consistent with other studies where nutrient application rates were lower (Cramer et al. 2010; Barbosa et al. 2014b). This suggests that N and P deposition, at rates expected by 2030, are unlikely to have pronounced short-term effects on tree recruitment in Indian savannas and tropical dry forests with respect to tree seedling growth rates and accumulation of ecosystem fuel loads.

N-fixers demonstrated very high levels of seedling survival that was unaffected by nutrient or fire treatments. Seedling mortality was almost exclusively restricted to non-N-fixers. Survival of non-N-fixers was significantly lower than N-fixers in the unburned treatment (Fig 4) and declined substantially following burning. In fact, only half of non-N-fixers survived fire, compared to 99% of N-fixers. These results reiterate the important role fire plays in restricting tree recruitment, and hence, tree cover in this ecosystem that numerous studies have previously highlighted (Bond and van Wilgen 1996; Higgins et al. 2000; Sankaran et al. 2005; Bond 2008). In addition, it also demonstrates the role of differential susceptibility to fire-mediated mortality between N-fixers and non-N-fixers in shaping tree community composition, an aspect that has thus far not been adequately addressed.

One reason for greater survival of N-fixers could be their faster growth rates compared to non-N-fixers, which allows them to reach larger seedling sizes before the start of the fire season, which in turn confers them with increased post-fire survival (Trollope 1984; Gignoux et al. 1997; Hoffmann and Solbrig 2003; Bond 2008; Lawes et al. 2011; Wakeling et al. 2011). Further, N-fixing tropical savanna and dry forest tree seedlings have been shown to partition relatively greater amounts of biomass to roots, as well as invest in higher root storage carbohydrate concentrations (Varma et al. 2018). This below-ground investment is positively associated with post-fire resprouting (Hoffmann et al. 2000; Bell 2001; Bond et al. 2003b; Bond and Midgley 2003; Clarke and Knox 2009; Wigley et al. 2009; Schutz et al. 2009; Clarke et al. 2013), which can promote survival in N-fixers.

Larger stem diameters were positively associated with post-fire survival probabilities amongst non-N-fixing species. With nutrient addition, and in particular N addition, non-N-fixers attained larger maximum stem diameters compared to individuals from control treatments, a result of consistent increases in stem diameter growth rates of the fastest growing individuals with N addition. On average, across the four non-N-fixing species, 16% of individuals had greater stem diameters in the N addition treatment compared to the maximum observed in the control treatment. Consequently, these individuals also had higher post-fire survival probabilities. As N-fixers displayed very high baseline rates of post-fire survival, they had no room for improved survival with increases in seedling size. Therefore, with nutrient deposition, the community of surviving seedlings or resprouts could see an increase in the relative abundance of non-N-fixers.

In addition to fire, herbivory constitutes another major disturbance agent in this ecosystem that can influence recruitment and composition of tree communities (Frost et al. 1986; Sankaran et al. 2004, 2008). Herbivory can interact with fires to impose a stronger control on tree cover, compared to that exerted by either disturbance agent independently (Staver et al. 2009). Susceptibility to herbivory, as well as plant investment into herbivory defence can be mediated by nutrient availability (Bryant et al. 1983; Endara and Coley 2011; Vadigi and Ward 2012). Hence, further research on the impacts of nutrient deposition on tropical dry forest tree recruitment would benefit from the explicit consideration of the interaction between nutrient deposition, fire regimes and herbivory. Additionally, in disturbance prone ecosystems, post-disturbance recovery of tree seedlings is a key process that influences tree recruitment, as it can play an important role in determining susceptibility of plants to subsequent disturbance events (Bond and Midgley 2001; Wigley et al. 2009; Schutz et al. 2009). Previous studies have shown that post-disturbance resprouts can differ substantially from seedlings in terms of growth (Hoffmann and Solbrig 2003; Grady and Hoffmann 2012). However, differences in patterns of recovery between species and functional groups, and the influence of nutrient availability is relatively unexplored.

Atmospheric nutrient deposition has the potential to alter vegetation communities through direct effects of fertilisation on growth and productivity of plants and indirectly, by modifying properties of disturbance regimes, as well as the susceptibility of plants to disturbance events. Quantifying these effects within the context of processes that shape vegetation communities is critical towards assessing the future trajectory of ecosystems. In this study we demonstrated that greater seedling growth rates and consequently, high post-fire survival in N-fixing species compared to non-N-fixers may play an important role in structuring tree communities in tropical dry forests. Rates of nutrient deposition in the near future are unlikely to result in large modifications in this baseline advantage for N-fixers in our study area. However, even with conservative rates of nutrient deposition, increased nutrient availability (especially N) has the potential to modify seedling growth rates. In non-N-fixers, this leads to increases in post-fire survival probabilities. As N-fixers maintain high levels of post-fire survival, that are unaffected by nutrient addition, the resprouting community may see increases in the relative representation of non-N-fixers, that in the long-term, could be reflected in adult tree assemblages.

## Acknowledgements

We would like to thank Kavita Isvaran, Suhel Quader and Jayashree Ratnam for their input during planning and execution of the experiment. We also thank Dr. Ganesh Babu and Umesh VJ at the Foundation for the Revitalisation of Local Health and Tradition (FRLHT), Bangalore, for providing the tree seedlings, and Manjunatha HC, Meenakshi HJ, and Chandregowda J for the use of their land for the experiment. Our gratitude to our field assistants Mahesh HK, Bomrai HK and Mahadev HK, and also to Hemanth Kumar HV, Anil PA, Harinandanan PV, Chengappa SK, Rutuja Dhamale, Mayank Kohli, Nandita Nataraj and Prashanth Gowda for their assistance during the experiment. Funding for this work was provided by the National Centre for Biological Sciences (NCBS), Bangalore. This manuscript was greatly improved by comments from Anand M Osuri, Fiona Savory, Sumanta Baghchi and Edmund C February.

